# Chromosome-level genome of the three-spot damselfish, *Dascyllus trimaculatus*

**DOI:** 10.1101/2022.08.16.504202

**Authors:** May B. Roberts, Darrin T. Schultz, Remy Gatins, Merly Escalona, Giacomo Bernardi

**Affiliations:** Department of Ecology and Evolutionary Biology, 115 McAllister Rd, University of California Santa Cruz, CA, USA; Department of Molecular Evolution and Development, University of Vienna, Vienna, 1010, Austria; Monterey Bay Aquarium Research Institute, Moss Landing, California 95039, USA; Department of Biomolecular Engineering and Bioinformatics, University of California, Santa Cruz, California 95064, USA; Department of Marine Sciences, Northeastern University, MA, USA

**Keywords:** Hybrid genome assembly, Robertsonian polymorphism, chromosome fusion, domino damselfish, ONT, Hi-C Chicago, Illumina shotgun, coral reef fish, Pomacentridae

## Abstract

Damselfishes (Family: Pomacentridae) are a group of ecologically important, primarily coral reef fishes that include over 400 species. Damselfishes have been used as model organisms to study recruitment (anemonefishes), the effects of ocean acidification (spiny damselfish), population structure and speciation (*Dascyllus*). The genus *Dascyllus* includes a group of small bodied species, and a complex of relatively larger bodied species, the *Dascyllus trimaculatus* species complex that comprises several species including *D. trimaculatus* itself. The three-spot damselfish, *D. trimaculatus* is a widespread and common coral reef fish species found across the tropical Indo-Pacific. Here we present the first genome assembly of this species. This assembly contains 910 Mb, 90% of the bases are in 24 chromosome-scale scaffolds, and the BUSCO score of the assembly is 97.9%. Our findings confirm previous reports of a karyotype of 2n = 47 in *D. trimaculatus* in which one parent contributes 24 chromosomes and the other 23. We find evidence that this karyotype is the result of a heterozygous Robertsonian fusion. We also find that the *D. trimaculatus* chromosomes are each homologous with single chromosomes of the closely related clownfish species, *Amphiprion percula*. This assembly will be a valuable resource in the population genomics and conservation of Damselfishes, and continued studies of the karyotypic diversity in this clade.

## Introduction

Damselfishes (Pomacentridae) are a group of small-bodied species found across all coral reef regions and most temperate marine systems where they are often the most visibly abundant fishes on the reef (1–4). This family includes more than 400 species that, despite their small size (max 30cm), play important ecological roles (2,5). Within this large family, the genus *Dascyllus* comprises eleven species, four of which, make up the *Dascyllus trimaculatus* species complex. This species complex includes three described species with restricted geographic ranges, *D. albisella* in the Hawaiian Islands, *D. strasburgi* in the Marquesas Islands, and *D. auripinnis* in the Line Islands. In contrast, *D. trimaculatus* has the broadest range, extending from the Red Sea, where it was first described (Ruppell, 1829), across the tropical and subtropical Indo-Pacific (Figure 1).

**Figure 1.**
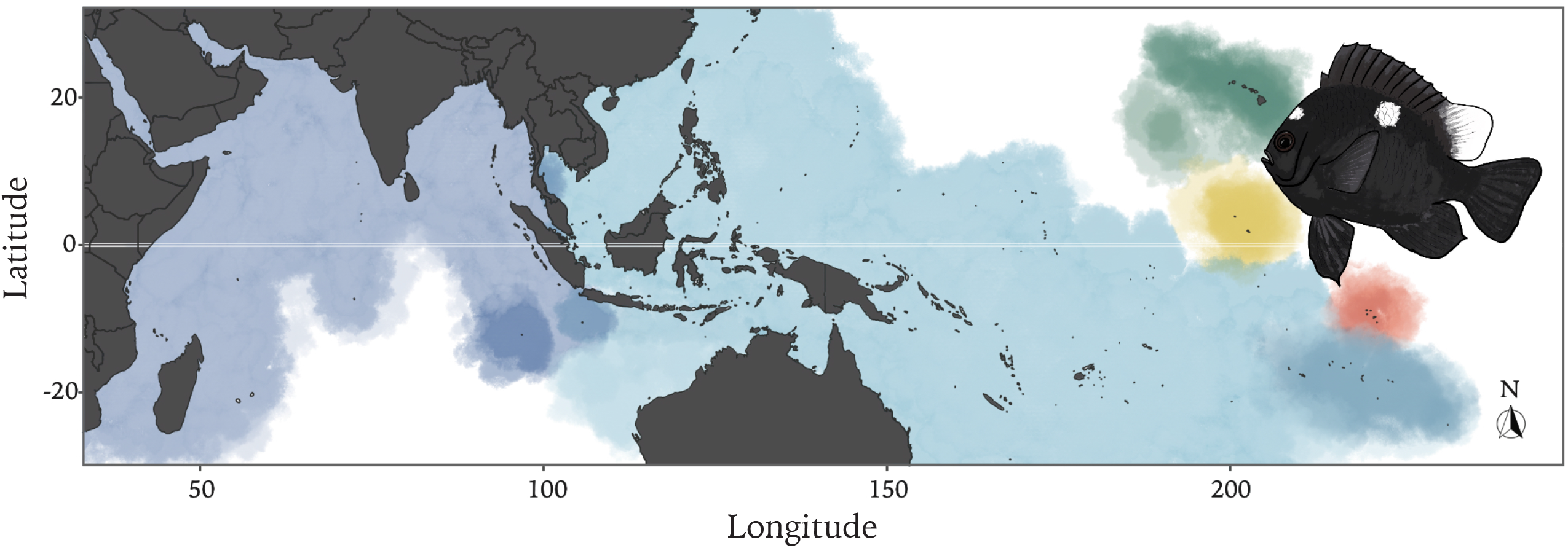
*Dascyllus trimaculatus* and species complex global distribution. The three-spot damselfish (Pacific morph) is shown on the right of the map adapted from Leray et al. (2009) and Salas et al. (2020). The map shows in blues the broad distribution of *D. trimaculatus*. Differences in blues reflect results of Salas et al. (2020) which showed Indian Ocean differentiation from Pacific populations as well as a sub population in Cocos Keeling and a hybridization zone in Christmas Island. Similarly, the darker blue patch in the Central Pacific shows another divergent population of *D. trimaculatus* identified in Leray et al (2009). The other colors show the distributions of the other species within the *D. trimaculatus* complex: green for *D. albisella*, yellow for *D. auripinnis*, and red for *D. strasburg*.

Three-spot damselfish is an abundant and common species, that exhibit a typical bipartite life history, with a site attached adult phase, where mate pairs lay, fertilize, and care for demersal eggs, followed by a pelagic larval phase. Larvae hatch after ∼6 days and feed in the water column on zooplankton where their pelagic larval duration (PLD) lasts 23-30 days until they recruit back to the reef (6,7). Larvae settle primarily into anemones for protection often sharing this shelter with different species of the popular anemonefish (in Hawai’i, where anemonefishes and anemones are absent, *D. albisella* recruits to branching coral). As subadults, they leave the anemone and live nearby in small to large groups.

There has also been considerable effort in understanding the chromosomes architecture and variation of *Dascyllus* and other damselfishes. Chromosome number varies between species of *Dascyllus* as well as within species (8–10) giving insight into chromosomal drivers of evolution (11–13) and how this variation is manifested ecologically (11,14). As we shift into the age of genomic natural history where genomic tools offer vastly more detail and statistical power, a reference genome will aid in further refining our understanding of wildlife biology (15). There are currently 14 Pomacentrid reference genomes, five of which are publicly available through the National Center for Biotechnology Information (NCBI; https://www.ncbi.nlm.nih.gov/) (*A. ocellaris*, (16), *Acanthochromis polyacanthus*, (17); *Amphiprion percula*, (18); *Amphiprion ocellaris* (19); *Acanthochromis polyacanthus*, Lehmann in review), and another nine from a single study (Marcionetti et al. 2019). Of these, only one is of species other than genus *Amphiprion* and only three of those listed above (*A. ocellaris, A. percula, A. polyacanthus*) are chromosome-scale genomes. Of the Pomacentrid chromosome-scale genomes all had 2n=48, with genome sizes ranging between 863Mb-956Mb. The two published genomes, *A. ocellaris*, (Ryu et al. 2022) and *A. percula* were highly complete with published BUSCO values of 97.01% and 97.2% respectively. Chromosome-scale genomes provide a more complete sequence and locations of genes and allow for research into how chromosome architecture influences ecology, population dynamics, and adaptive evolution. Here, we present the first genome assembly within the genus *Dascyllus* and add to the short, but growing list of Pomacentrid chromosome-scale genomes.

## Materials and Methods

### Biological Materials

The *D. trimaculatus* individual used for this genome assembly was ordered from an online pet fish supplier (liveaquaria.com), sourced from the West Pacific Rim population (20). It was euthanized following an approved IACUC protocol animal use. Liver, muscle, gill, and brain tissue were harvested from the right side of the individual and each placed in separate, pre-weighed Covaris cryogenic vials, flash frozen in liquid nitrogen, and stored at -80°C until further processing. The remaining intact left-side of the specimen is stored in -80°C at University of California Santa Cruz.

### Nucleic acid library preparation and sequencing

#### Whole-genome shotgun library preparation

DNA was extracted from 13 mg of muscle tissue using a DNeasy Blood and Tissue kit (Qiagen), quantified using Qubit dsDNA HS Assay kit (Thermo Fisher Scientific) and Qubit 4.0 Fluorometer, then assayed with 1.0% agarose gel electrophoresis to determine molecular weight. DNA was sheared for 26 cycles of shearing (15s on, 30s chilling) using a Bioruptor sonicator (Diagenode), then size selected using SPRI beads (Beckman) to select for fragments between 200 bp to 500 bp.

The NEBNext UltraII DNA Library Prep Kit for Illumina (New England Bio Labs) was used according to manufacturer’s protocol except that KAPA Hot Mix Ready Start Master Mix (Roche Diagnostics) was used for library amplification instead of NEB Q5 Master Mix. Paired-end sequencing was done at the University of California Davis Genome Center on a HiSeq4000 sequencer on a 2×150PE cycle.

#### Chicago library preparation

High molecular weight (HMW) DNA was isolated from the *Dascyllus trimaculatus* individual by lysing gill tissue in low-EDTA TE buffer (21), then purifying with a chloroform, phenol:chloroform, chloroform, ethanol precipitation protocol (22). The quality of the HMW DNA was assayed with 1.0% agarose gel electrophoresis. This DNA was used in the preparation of the Chicago, Hi-C, and for Oxford Nanopore Technologies sequencing libraries.

From this DNA, three Chicago libraries were prepared using a published method (23), each using a different restriction enzyme: one with DpnII cutting at GATC sites, one with MluCI cutting at AATT sites, and one with FatI cutting at CATG sites. These libraries were sequenced on a 2×150PE cycle at Fulgent Genetics on a HiSeq400 sequencer.

#### Hi-C library preparation

Two Hi-C libraries were generated from approximately 100 ng of LN^2^-flash-frozen muscle. The libraries were constructed using a published protocol (24). One library was constructed using the enzyme DpnII, and the other library was constructed with the enzyme MluCI.

#### Oxford Nanopore library (ONT)

Next, 1500 ng of the HMW DNA prepared for Chicago libraries was also used to prepare two ONT WGS libraries with the SQK-LSK109 modified protocol “versionGDE_9063_v109_revT_14Aug2019”. The DNA repair steps at 20°C and 65°C were carried out for 20 minutes each, instead of 5 minutes each. We ran each of the resulting libraries on two separate MinION flow cells (FLO-MIN106), each for 72 hours. Raw fast5 files from the two MinION runs were basecalled using Guppy(25) v3.3.

A summary of sequencing information for the various libraries can be found in Supplementary Table 1.

### Genome Assembly

All programs and versions used for the assembly are listed in Table 1. Sequencing adapters were removed from the Illumina whole-genome shotgun (WGS) reads with Trimmomatic (26) v0.39 with parameters: ‘ILLUMINACLIP: all_seqs.fa:2:30:10:8:TRUE SLIDINGWINDOW:4:20 MINLEN:50’. We used jellyfish (26) v2.2.10 to make a k-21 k-mer count vs abundance histogram and used the histogram with Genome Scope (27) v2.0 to estimate *D. trimaculatus* genome size, heterozygosity, and repeat content. MaSuRCA was used to assemble a first version of the genome using both the ONT and WGS reads.

**Table 1.** List of programs and program versions in order of use for the genome assembly of the three-spot damselfish, *Dascyllus trimaculatus*

| Purpose |  | Software and version |
| --- | --- | --- |
| Estimate genome size |  | Jellyfish v2.2.10 |
| Reference-free characterization |  | GenomeScope v2 |
| Basecalling |  | Guppy v3.3 |
| *De novo* assembly |  | MaSuRCA downloaded Sept 2020 |
| Map HiC/Chicago reads |  | Arima HiC pipeline Release date: 05. 2019<br>BWA v0.7.17-r1188; samtools v1.13; Picard v2.26.2 |
| File processing |  | Samtools v1.13, bedtools v2.30.0 |
| Proximity ligation scaffolding |  | SALSA2 v2.3 |
| Trim adapters and filter |  | Trimmomatic v0.39 |
| Polish assembly (2x) |  | Pilon v1.23 |
| Check for contamination |  | Blobtools2 v3.1.0 |
| Manual curation | Map Hi-C to genome | Chromap v0.2.2 |
|  | File conversion | Juicebox Assembly Tools v2.14.0 |
|  |  | Artisinal tools 9a79889<br>https://github.com/conchoecia/genome_assembly_pipelines b0cda60 |
|  | Chromosome assignment | D-Genies Accessed April, 2022 |
| Assembly statistics | Completeness | BUSCO v5.2.2, assembly-stats v0.0.1, merqury v1.3 |
|  | General stats | Assembly-stats v0.0.1 |
|  | Quality and error rate | Merqury v1.3 |

We followed the Arima-HiC mapping pipeline (https://github.com/ArimaGenomics/mapping_pipeline/blob/master/Arima_Mapping_UserGuide_A160156_v02.pdf) to prepare the data for scaffolding. The pipeline aligns the sequencing data from each the Hi-C and Chicago dataset against the assembly from MaSuRCA, it then filters ligation adapters and removes PCR duplicates from the resulting alignments. These alignments were then processed with samtools (31,32) v1.13 and converted into BED files with bedtools (33) v2.30.

The MaSuRCA assembly was scaffolded with SALSA (34) v2.3 with ligation junction parameter -e AATT,GATC,CATG. Iteration number was set to 10 (-i 10) and we allowed for Hi-C/Chicago data to also correct assembly errors (-m yes).

We aligned the trimmed Illumina WGS reads to the scaffolded output of SALSA with bwa mem (35) v0.7.17-r1188 and used that alignment to polish the assembly with Pilon (36) v1.23. We repeated the alignment and polishing steps once. The error-corrected assembly was then screened for possible contaminants, using Blobtools2 (37) v3.1.0. Any contigs assigned to phyla other than Chordata were removed. However, any sequences categorized as ‘No hits’ were kept. The assembly was then manually curated by mapping the DpnII and MluCI Hi-C reads to the genome assembly with chromap (38) v0.2.2 with a quality filter of 0 and converted to a .hic file with Juicebox Assembly Tools (JBAT) (39) v2.14.00. Artisanal tools commit 9a79889 (https://bitbucket.org/bredeson/artisanal) was used to generate a JBAT assembly file. We used the Juicebox GUI (40) v1.11.08 to manually curate the assembly with the .hic and .assembly files. Modifications made to the assembly included ordering and orienting scaffolds into chromosome-scale scaffolds, removing duplicated regions, and making manual assembly breaks to place misassembled contig pieces onto the correct scaffold. Artisanal was used to generate an updated genome assembly FASTA file. Scaffolds not placed on chromosomes were sorted by the strongest Hi-C connection to chromosome-scale scaffolds with genome assembly tools commit b0cda60 (https://github.com/conchoecia/genome_assembly_pipelines. D-Genies (41), accessed April 30th 2022, was used to align the manually-curated assembly to the chromosome-scale assembly of the closely related *Amphiprion percula* genome assembly (18). The evidence from this analysis was used to assign chromosome numbers to the *D. trimaculatus* scaffolds based on homology with *A. percula* chromosomes.

### Genome quality assessment

Benchmarking Universal Single-Copy Orthologs (BUSCO) (42,43) v5.2.2 was used to evaluate genome completeness by comparing number of orthologous genes found in the assembly to the 3,640 genes in the actinopterygii_odb10 database. Assembly statistics (assembly-stats; https://github.com/sanger-pathogens/assembly-stats) were generated to track N50, L50, contigs, gaps, and lengths at each step. We used merqury (44) v1.3, to calculate the genome completeness and error rates.

## Results

### Sequencing

We sequenced four library types: a whole-genome shotgun library which resulted in 314.6Mb paired-end 150bp reads, representing 103x coverage, and 3.52M (4.84Gb) and 8.57M (19.77Gb) ONT reads from the two runs on the minION flowcells, representing 22x ONT coverage for the initial hybrid assembly. The five proximity ligation libraries used for scaffolding, two Hi-C (restriction enzymes DpnII and Mlucl), and three Chicago libraries (restriction enzymes DpnII, MlucI, and FatI) yielded ∼108M, ∼152M, ∼65M, ∼74M, ∼67M, reads respectively for a total proximity ligation coverage of 154x. In total, across all data types, we had a final coverage of 280x (See Supplementary Table 1 for sequencing details).

### Heterozygosity and repetitive sequence estimation

GenomeScope estimated the genome size to be 809 Mb, with 84% unique and 16% repetitive sequences, and 1.02% heterozygosity (Supplementary Figure 1).

### Genome Assembly

Genome quality metrics for each step of the assembly are listed in Table 2. The initial *de novo* assembly by MaSuRCA with ONT and Illumina shotgun data had a total length of 919,275,268bp in 3,501 contigs with an N50 of 1,108Kb. Scaffolding with the HiC and Chicago libraries dropped the number of contigs to 2,467 and increased N50 to 16,013Kb. After two rounds of polishing with trimmed Illumina shotgun reads gaps decreased from 1,097 to 1,088. Blobtools2 showed that of the 2,467 contigs, none matched other taxa in NCBI databases of bacteria, invertebrates, mammals, phages, plants and fungi, or environmental samples. Four hundred seventy-eight contigs did not match any databases (‘no-hits’) and were left in the genome.

**Table 2.**
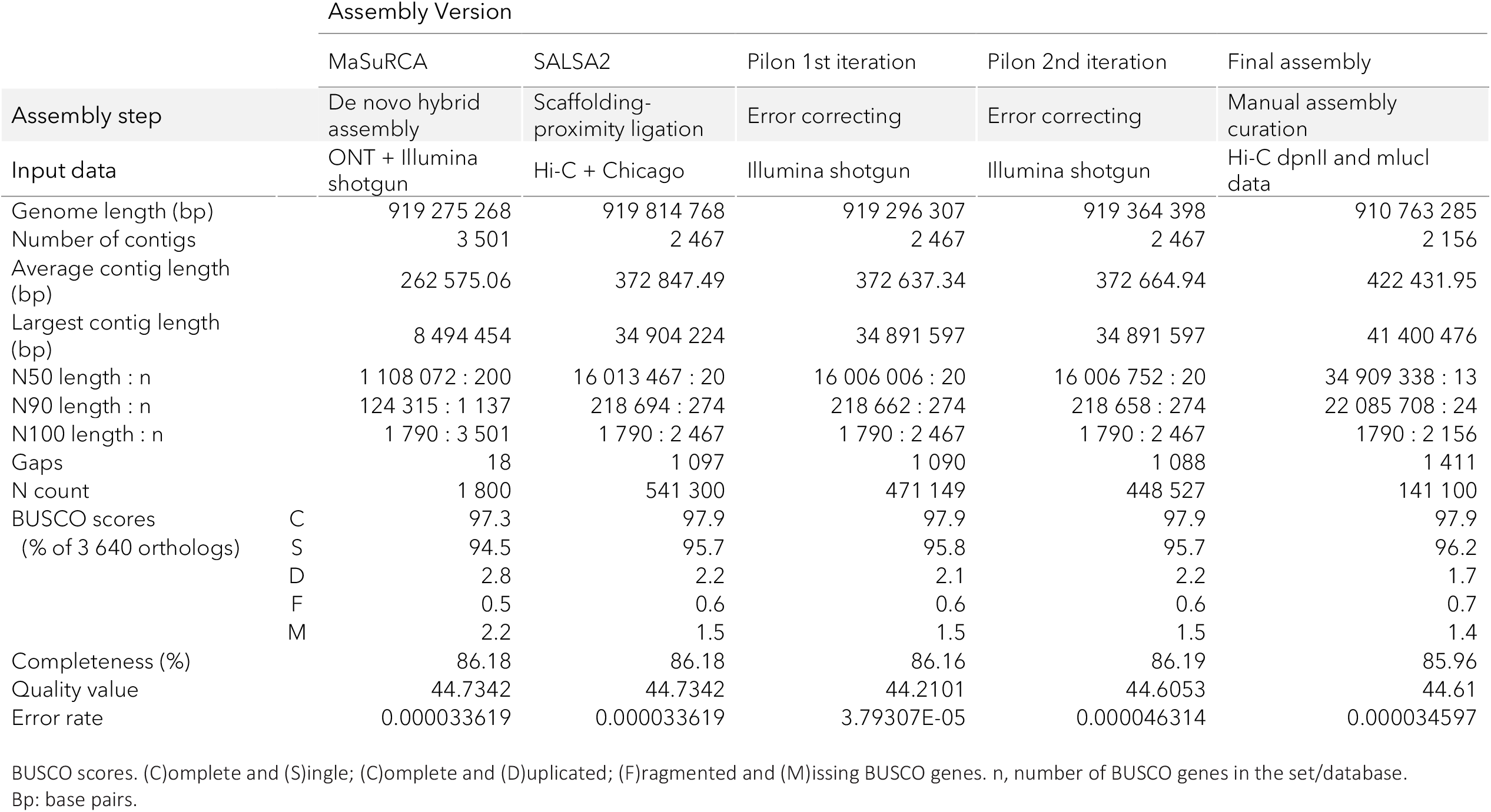
A comparison of genome metrics between *D*.*trimaculatus* assembly stages.

The manual curation of the genome assembly yielded 24 scaffolds consistent with chromosome-scale scaffolds (Figure 2). A dotplot comparison (Figure 3) with the *Amphiprion percula* (18) genome revealed that each of the *D. trimaculatus* chromosome-scale scaffolds had a one-to-one corresponding homologous, albeit rearranged, chromosome in the *Amphiprion percula* genome.

**Figure 2.**
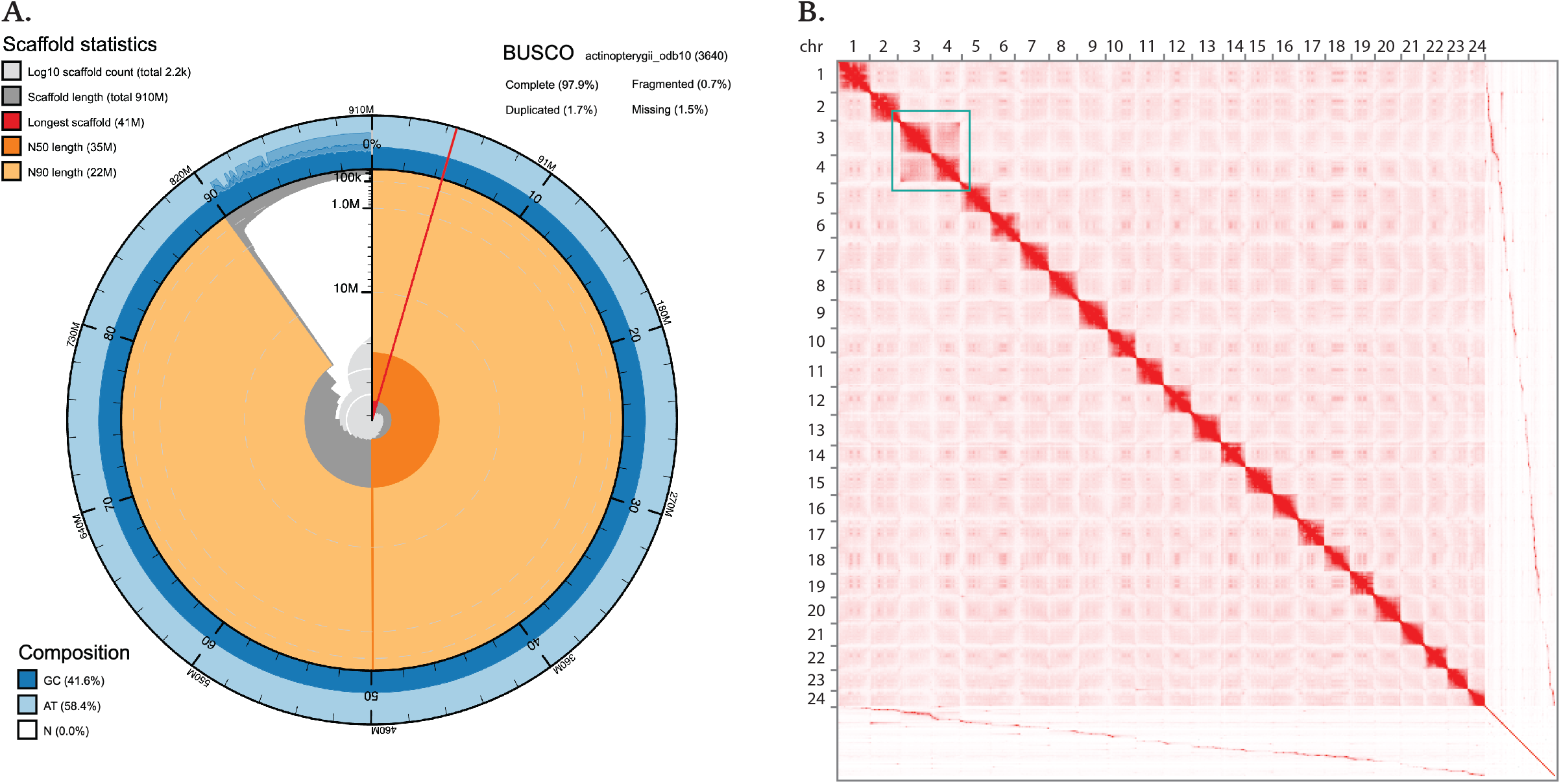
Genome statistics and chromosome map.Panel. **A:** The outer circumference of the main plot represents the full length of the 910,763,285 bp chromosome-scale assembly of *Dascyllus trimaculatus*. The outer ring of blues depicts GC (dark blue) and AT (light blue) content along the assembly which is summarized in the lower left. The second ring is demarcated by percentage of the total contigs of the genome. Orange and pale-orange arcs show the N50 and N90 record lengths (34,909,338 and 22,085,708 bp), respectively overlying the dark grey, which arranges scaffolds in order by size starting from the largest scaffold (41,400,476 bp and ∼4% genome, shown in red). The pale grey spiral shows the cumulative scaffold count on a log scale with white scale lines showing successive orders of magnitude. A summary of BUSCO statistics for complete (97.9%), fragmented (0.7%), duplicated (1.7%), and missing (1.5%), orthologous genes in the actinopterygii_odb10 set is shown in the top right. **Panel B:** A Hi-C contact map made with the MluCI and the DpnII libraries showing 24 chromosome clusters and the unscaffolded contigs. In the green square, chromosomes 3 and 4 show strong interchromosomal connections at roughly half coverage indicating Robertsonian fusion in one set of chromosomes contributed a parent with 2n=47 while the other parent contributed 2n=48.

**Figure 3.**
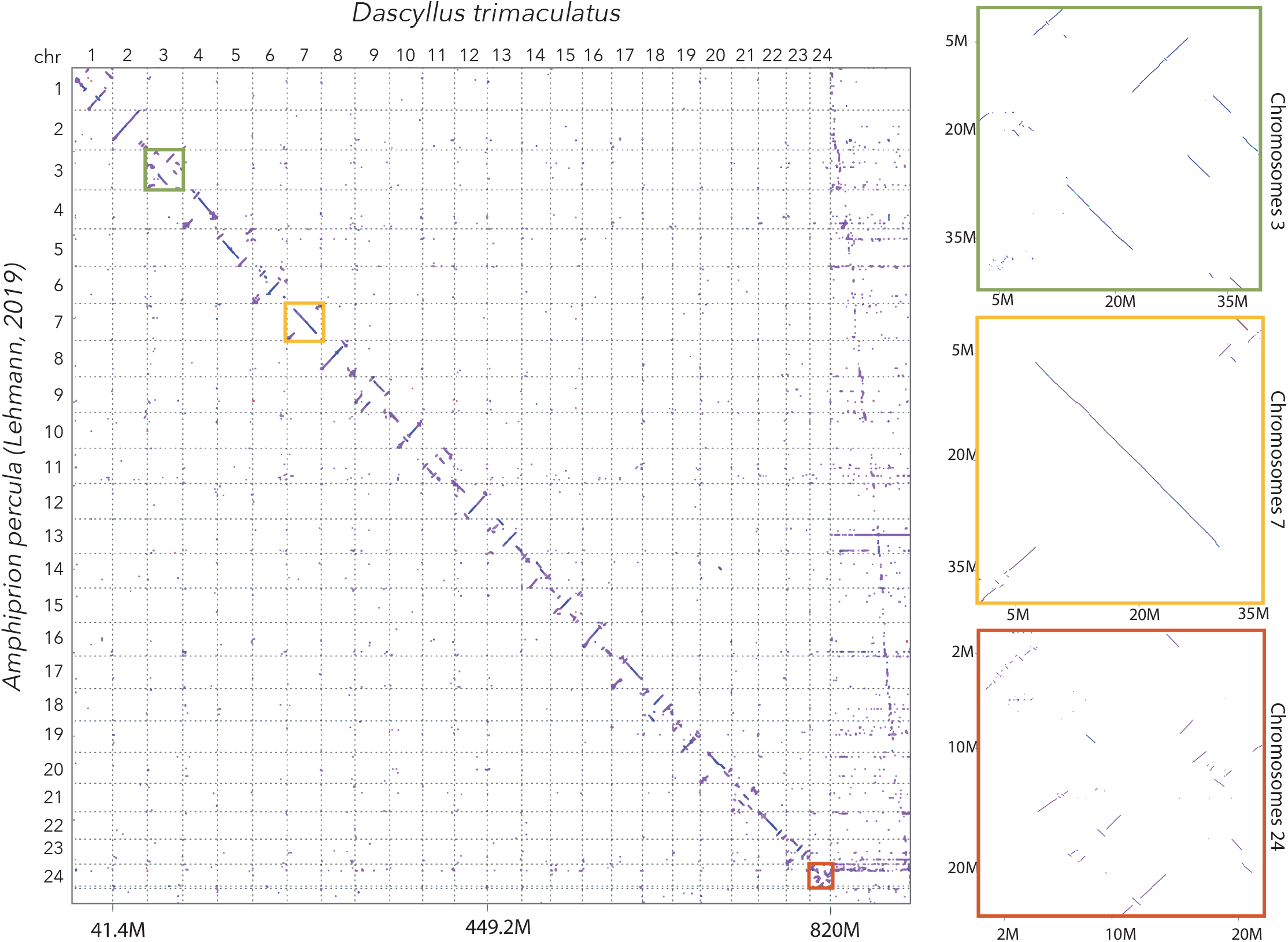
*Dascyllus trimaculatus* mapped against chromosome-level genome of *Amphiprion percula* (Pomacentridae) The main panel shows a dot-plot of the assemblies of *Dascyllus trimaculatus* (presented in this manuscript) and the genome of the anemone fish *Amphiprion percula* (Lehmann, 2019). The remaining un-scaffolded contigs are shown in the last column. The right three panels show a close-up dot plot of the color-coded boxes in the main panel of chromosome 3, chromosome 7, and chromosome 24. Chromosome 3, one of the chromosomes involved in Robertsonian fusions within the species, shows many rearrangements as well as regions of repeat sequences near one end. Chromosome 7 seems to be one of the most architecturally conserved chromosomes between *Dascyllu*s and *Amphiprion*, whereas Chromosome 24 shows and examples of a highly rearranged chromosome.

The final assembly (GenBank accession: JAMOIN000000000) has a length of 910.7Mb, 90% of which was on chromosome-scale scaffolds and BUSCO score of 97.9%. Merqury calculated 86.19% completeness, QV of 44.6, and an estimated error rate 0.0000346, or a single nucleotide error every 28.9 Kb.

## Discussion

The biology, evolution and biogeography of the three-spot damselfish is relatively well-studied using genetic (10,45–51) and genomic tools (52,53) and, as we shift further into the age of WGS data and tools, a reference genome is an invaluable resource. Here we present the chromosome-scale genome assembly of a three-spot damselfish, *Dascyllus trimaculatus*, collected from the Indonesian/Philippine population (20). It is the first within the genus *Dascyllus* of the widely studied, and large Pomacentridae family. This high-quality *de novo* assembly of a non-model coral reef fish is a valuable reference for furthering studies of evolutionary, ecological, and conservation studies for the species and for coral reef fish in general.

We report sequences for 24 chromosomes of the *D. trimaculatus* genome with total length and repetitive content (Figure 2A, Supplementary Figure 1) that is expected for this species (10,54,55). Interestingly, our Hi-C data also show that chromosomes three and four have strong connections at half the depth of other intra-chromosomal connections (Figure 2B). This pattern can be explained by a hemizygous state wherein one parental gamete contributed a Robertsonian-fusion of chromosomes three and four, and the other parental gamete contributed chromosomes three and four as separate chromosomes making the individual sequenced here, a 2n=47 individual. This finding is consistent with previous studies that report both 2n=47 and 2n=48 for *Dascyllus trimaculatus* (8,9,54). Chromosome numbers vary both within and among species of *Dascyllus*. One report on several *Dascyllus* species collected in the Philippines and the Ryukyu Archipelago of southern Japan demonstrated polymorphic karyotypes in all but one of the species (Ojima and Kashiwagi, 1981). *Dascyllus aruanus* had the most karyotypic variation - between 2n = 27-33 chromosomes, *D. reticulatus* 2n = 34-37, *D. trimaculatus* 2n = 47-48, and *D. melanurus* with 2n = 48.

In addition to confirming variation in chromosome number, the dot plot comparison between this genome and of the closest relative with an available chromosome-scale assembly, *Amphiprion percula* (18), revealed several rearrangements in every chromosome between corresponding chromosomes (Figure 3). The Pomacentrid subfamilies Chrominae and Amphiprionini are estimated to have diverged over 50 million years ago (mya) (56). The estimated number of rearrangements within chromosomes ranged from 2+ in chromosome 7 of *D. trimaculatus* which was the most like its counterpart in *A. percula* to over 35 in chromosome 24 (Figure 3). This pattern of rearrangements has not been characterized between chromosome-scale genome assemblies of Pomacentridae. The role of variation in chromosome number has been the subject of several cytogenic studies which have found that chromosome diversity inversely related to mobility of the fish and that chromosome rearrangements can serve to either promote or prevent recombination events (11,12,14,57). Interestingly, chromosome 3 in the genome presented in this paper is one of the most rearranged while also being one of the chromosomes involved in the Robertsonian fusion mentioned above. This assembly will be a useful starting point to study how this type of genome structure varies at a meta-population scale, and how this influences recombination and adaptation.

This assembly represents the first chromosome-level genome of the genus *Dascyllus* as well as the first non-*Amphiprion* chromosome-scale genome published in the Pomacentridae family. Damselfishes are excellent model species due to their relatively small size, ease to manage in the wild and lab, and those interested in this group will benefit from this addition to the available genomic resources. *Dascyllus trimaculatus* itself, is has had a dynamic evolutionary trajectory across the Indo-Pacific, evident in species complex that is continuing to reveal its complexity and provide insight into evolutionary mechanisms. In addition to providing a high-quality reference genome to further our understanding of genomic architecture, this assembly will serve to leverage information stored across the genome to better understand the population dynamics, phylogeny, biogeography, demographics, of *Dascyllus trimaculatus*, as well as gain insight into historical, current, and future response to changes in climate.

## Data Availability

The assembly and genomic sequencing reads generated for this study have all been deposited in the NCBI GenBank database under BioProject ID PRJNA828170. The accession for the genome is JAMOIN000000000, WGS data (XX), proximity ligation data (XX, XX, XX, XX, XX), ONT data (XX, XX).

## Acknowledgements

The authors are grateful for access to the Hummingbird computational cluster and the team behind it at University of California Santa Cruz. The whole genome shotgun sequencing was carried out at the DNA Technologies and Expression Analysis Core at the University of California Davis Genome Center, supported by NIH Shared Instrumentation Grant 1S10OD010786-01. We also appreciate Jonas Oppenheimer and Robert Lehmann for helpful discussions and advice.

## Figures

**Figure S1. GenomeScope results from paired-end 150bp whole genome sequences of *Dascyllus trimaculatus*.**

**Table S1.**
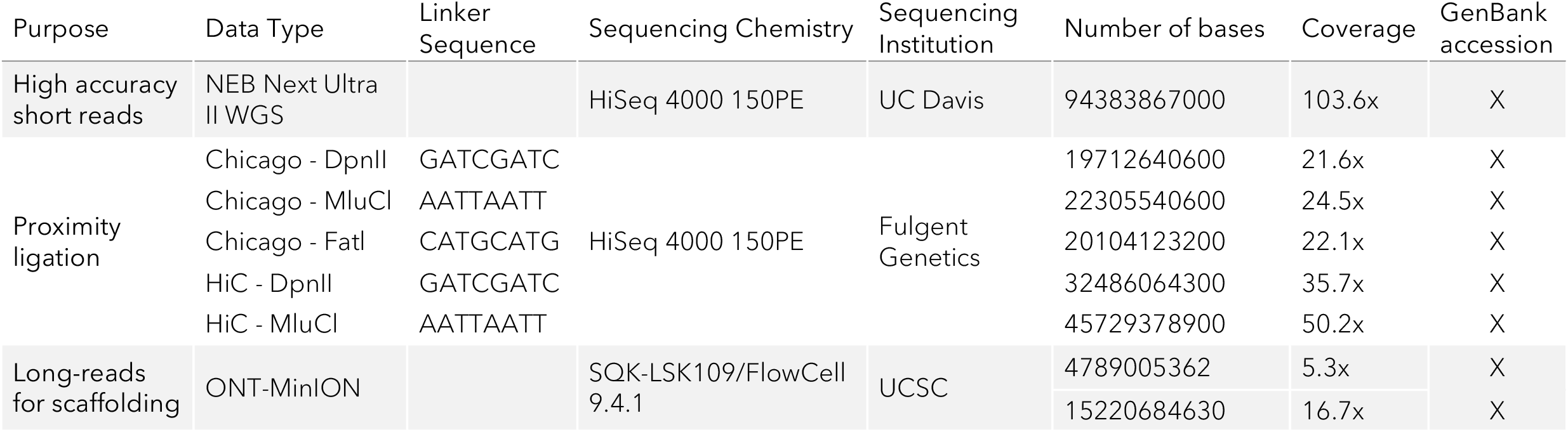
Sequencing information of data used to assemble the genome of the three-spot damselfish, *Dascyllus trimaculatus*, Bioproject: PRJNA828170 Biosample: SAMN27642109; Genome accession: JAMOIN000000000 for isolate Kuro_0920G

